# Assessing species coverage and assembly quality of rapidly accumulating sequenced genomes

**DOI:** 10.1101/2021.10.15.464561

**Authors:** Romain Feron, Robert M. Waterhouse

## Abstract

Ambitious initiatives to coordinate genome sequencing of Earth’s biodiversity mean that the accumulation of genomic data is growing rapidly. In addition to cataloguing biodiversity, these data provide the basis for understanding biological function and evolution. Accurate and complete genome assemblies offer a comprehensive and reliable foundation upon which to advance our understanding of organismal biology at genetic, species, and ecosystem levels. However, ever-changing sequencing technologies and analysis methods mean that available data are often heterogeneous in quality. In order to guide forthcoming genome generation efforts and promote efficient prioritisation of resources, it is thus essential to define and monitor taxonomic coverage and quality of the data. Here we present an automated analysis workflow that surveys genome assemblies from the United States National Center for Biotechnology Information (NCBI), assesses their completeness using the relevant Benchmarking Universal Single-Copy Orthologue (BUSCO) datasets, and collates the results into an interactively browsable resource. We apply our workflow to produce a community resource of available assemblies from the phylum Arthropoda, the Arthropoda Assembly Assessment Catalogue. Using this resource, we survey current taxonomic coverage and assembly quality at the NCBI, we examine how key assembly metrics relate to gene content completeness, and we compare results from using different BUSCO lineage datasets. These results demonstrate how the workflow can be used to build a community resource that enables large-scale assessments to survey species coverage and data quality of available genome assemblies, and to guide prioritisations for ongoing and future sampling, sequencing, and genome generation initiatives.

## Introduction

Advances in sequencing technologies are bringing down costs and reducing sample requirements, leading to an accelerating accumulation of new and improved genome assemblies. Ambitious initiatives to coordinate sequencing of all known species are generating representative genomes from across the tree of life that catalogue Earth’s genetic biodiversity. In addition to constituting an inventory of biological diversity, the assembled and annotated genomes drive research to understand function and evolution at multiple levels, as well as to benefit human welfare (Richards 2015; Lewin et al. 2018). Investigating such questions using genomic data often requires comprehensive multi-species comparative analyses that benefit from high quality assemblies (Zoonomia Consortium 2020; Feng et al. 2020). It is therefore essential to be able to define the current taxonomic coverage of high-quality assemblies in order to guide forthcoming sequencing efforts and promote efficient prioritisation of resources globally.

Methods to gauge assembly quality include two main families of metrics (Thrash et al. 2020). One summarises contiguity using metrics like N50 length, where half the assembly comprises sequences of length N50 or longer, or L50 count, the smallest number of sequences whose lengths sum to 50% of the assembly. Complementary approaches estimate completeness by examining gene or protein content, e.g. the DOmain-based General Measure for transcriptome and proteome quality Assessment, DOGMA (Dohmen et al. 2016; Kemena et al. 2019), or the Benchmarking Universal Single-Copy Orthologues, BUSCO (Simão et al. 2015; Waterhouse et al. 2018). BUSCO has emerged as a standard and is used by UniProt (The UniProt Consortium 2019) and the United States National Center for Biotechnology Information (NCBI) (Sayers et al. 2021), as well as by genomics data quality assessment pipelines like MultiQC (Ewels et al. 2016) and BlobToolKit (Challis et al. 2020). BUSCO is based on the evolutionary expectation that single-copy orthologues found in nearly all species from a given taxon should be present and single-copy in any newly sequenced species from the same clade. BUSCO datasets are built for multiple taxonomic lineages by identifying near-universal groups of single-copy orthologues from OrthoDB (Waterhouse et al. 2013; Zdobnov et al. 2021). For assembly evaluations, sequence searches followed by gene predictions and orthology classifications identify complete, duplicated, or fragmented BUSCOs. The proportions recovered indicate the completeness in terms of expected subsets of evolutionarily conserved genes. Extrapolating from these, a high BUSCO completeness score suggests that the sequencing and assembly procedure has successfully reconstructed a reliable representation of the full set of genes.

Using their Complete Proteome Detector algorithm, UniProt classifies available proteomes as ‘standard’, ‘close to standard’, or ‘outlier’, and more recently also provides BUSCO proteome completeness summaries. For assemblies, the NCBI Assembly database provides summary statistics and metadata for each record. Querying these can provide snapshots of taxonomic coverage and data quality, but researchers currently lack access to comprehensive and standardised assessments of the species coverage and quality of publicly available assemblies. Such resources would allow data producers to readily compare their assemblies to existing data at the most relevant taxonomic level. They would also provide researchers -from individual research groups, collaborations, consortia, to broader communities-with comprehensive coverage and quality overviews of available genome resources for their taxa of interest. Such communities would benefit from being able to regularly survey in a standardised manner the coverage and quality of available genomic resources for selected groups of species from their field of interest. This would (i) aid project design, particularly in the context of comparative genomics analyses; (ii) simplify comparisons of the quality of their own data with that of existing assemblies; and (iii) provide a means to survey accumulating genomics resources of interest to their ongoing research projects.

To address these needs, we developed an automated analysis workflow that performs BUSCO assessments of assemblies for user-selected taxa from the NCBI, concurrently collating assembly metadata to build a catalogue of metrics in a taxonomically-aware framework. To demonstrate the utility of standardised evaluations for a clade, we applied our workflow to the phylum Arthropoda, for which genome data are supporting research on a wide range of topics including their roles as pests and disease vectors (Childers et al. 2021). Since sequencing the fruit fly genome (Adams 2000), sampling of arthropods has included ants and other Hymenoptera (Favreau et al. 2018; Branstetter et al. 2018), arachnids (Garb et al. 2018), beetles (McKenna 2018), butterflies and other Lepidoptera (Triant et al. 2018), flies and other Diptera (Wiegmann & Richards 2018; Ruzzante et al. 2019), hemipterans (Panfilio & Angelini 2018), and many others (González et al. 2018; Richards et al. 2018). Through efforts such as the i5k 5000 arthropod genomes initiative (i5K Consortium 2013) and others, the arthropod genomics community has worked to overcome challenges in genome sequencing, assembly, and annotation (Brown & Tagu 2015; Waterhouse 2015; Li et al. 2019). Despite encompassing only a tiny fraction of all arthropod diversity and showing taxonomic biases in sampling, assemblies are accumulating rapidly and are now publicly available for hundreds of species (Hotaling et al. 2020, 2021).

Our large-scale assessments allowed us to (i) survey the current taxonomic coverage and assembly quality across Arthropoda; (ii) examine how key assembly metrics relate to gene content completeness; (iii) quantify effects on assessment resolution using different BUSCO lineage datasets; (iv) compare the results of BUSCO v3 with the newer BUSCO v4, and (v) demonstrate how our workflow can be used to build a community resource. We provide the catalogue as an open resource for the arthropod genomics community, and the standalone workflow for users to build their own catalogues tailored to the needs of their research communities. Enabling user-customisable, taxonomically-aware, standardised, and updatable quality assessments of available genome assemblies will empower genomics data producers and users, as well as helping to prioritise species for genomic sequencing of Earth’s biodiversity.

## Results and Discussion

### An automated workflow for assembly assessments

We developed an automated analysis workflow to build and maintain NCBI genome assembly assessment catalogues for selected taxa. This workflow performs the following steps: 1) query the NCBI GenBank Assembly database (Sayers et al. 2021) to retrieve information about available assemblies and corresponding metadata for a user-defined taxonomic group; 2) identify all relevant BUSCO lineages based on species taxonomy for each assembly; 3) run BUSCO on each assembly using each relevant lineage dataset; 4) generate a summary table that collates all BUSCO results with assembly metrics and metadata; and 5) generate an HTML / JavaScript interactive table containing all data from the summary (Figure S1). Assembly metadata are integrated into a summary file along with five metrics obtained from the results of running BUSCO on each assembly with each relevant lineage: the percentages of complete, complete single-copy, complete duplicated, fragmented, and missing BUSCOs. The workflow allows users to systematically assess all assemblies available at the NCBI for a given taxon of interest. Importantly, it is also designed to perform on-demand updates to assess assemblies added to NCBI GenBank since the last run. The final output provides all the information retrieved for each assembly in both JSON and tab-separated formats, and an HTML / JavaScript table is generated to display the data. This output is saved in a summary folder each time the workflow is run. The workflow is implemented using the Snakemake workflow management engine (Köster & Rahmann 2012; Mölder et al. 2021) and all software dependencies are managed by the Conda package manager. It is fully automated and can be configured using a yaml file to specify the query to use for the NCBI Assembly database, BUSCO parameters, and the information to display in the output tables. The code and documentation are available from https://gitlab.com/evogenlab/a3cat-workflow.

### A survey of arthropod genome assembly resources

Applying the assembly assessment workflow to the phylum Arthropoda on June 11, 2021 resulted in the retrieval of a total of 2083 assemblies from 1387 species, providing a snapshot of the taxonomic coverage of available genome resources for arthropods at the NCBI. Of the ∼120 arthropod orders recognised by the NCBI Taxonomy database (Schoch et al. 2020) or the Catalogue of Life (Roskov et al. 2020), 48 are represented by at least one genome assembly, with 21 orders represented by five or more assemblies (Figure 1). Currently available genome resources include 1929 assemblies for 1262 insect species and a further 154 assemblies for 125 other arthropod species. For Insecta, this is a doubling of the number of species since a November 2020 survey from (Hotaling et al. 2021). Species with assemblies represent a ∼0.06% sampling from a total of about 1 million described arthropod species (792,339 species records and 121 orders in the NCBI Taxonomy database on August 10, 2021; 1,126,288 extant species and 123 orders in the Catalogue of Life 2021-06-10 edition).

**Figure 1.**
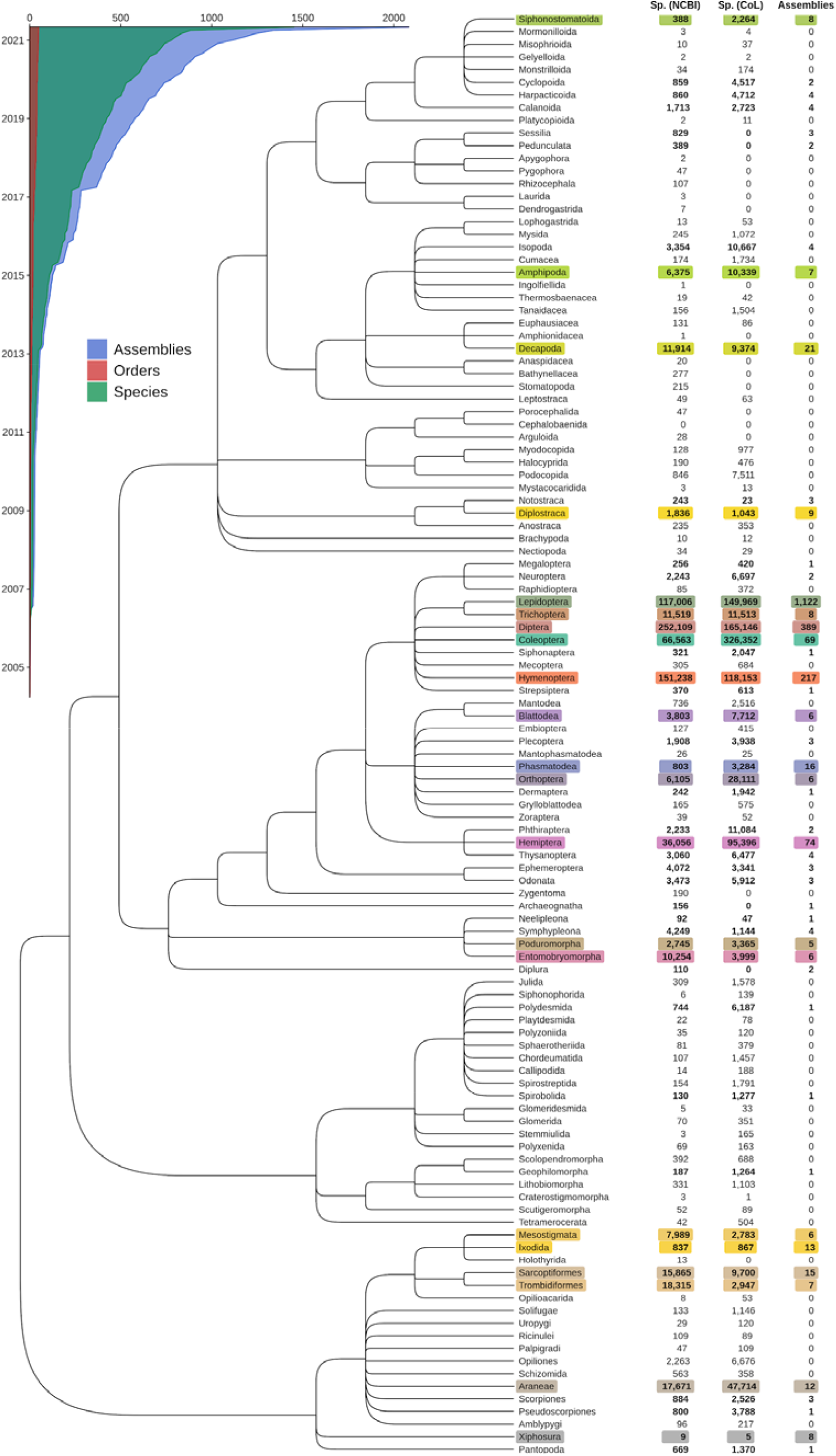
Available genome assembly resources across the arthropod phylogeny. The Arthropoda phylogeny from the United States National Center for Biotechnology Information (NCBI) Taxonomy database shows the evolutionary relationships amongst 114 orders. Counts of described species (Sp.) within each order are shown from the NCBI (v.2021-06-11) and the Catalogue of Life (CoL, v.2021-06-10), alongside numbers of genome assemblies available from the NCBI Assembly database (accessed on 2021-08-25). Of the 114 orders recognised by both the NCBI and the CoL, 48 orders are represented by at least one genome assembly. The 21 orders with five or more assemblies are highlighted with distinct colours, which are maintained for cross-referencing in Figures 2, 3, and 4. The inset shows the accumulation of assemblies, species, and orders submitted to the NCBI since 2005 (note that in the case of assembly updates only the latest submission dates are considered).

This survey highlights the sparsity and taxonomic imbalance of current species sampling, with 79.5% of species (83% of assemblies) belonging to only three orders: Lepidoptera -butterflies, moths, etc. (712 species, 1122 assemblies), Diptera - flies, etc. (216 species, 389 assemblies), and Hymenoptera - ants, bees, wasps, etc. (175 species, 217 assemblies). Similar sampling biases were identified by the November 2020 survey of NCBI resources for Insecta (Hotaling et al. 2021), where order-level counts for 601 insect species from 20 orders were 28% Diptera, 20% Lepidoptera, and 27% Hymenoptera. Notably, while about a third of insect orders are represented, only 5 to 10% of orders from other groups such as crustaceans, myriapods (centipedes, millipedes, etc.), and chelicerates (spiders, scorpions, etc.) have at least one assembly. Across Arthropoda, orders with the most sequenced species also show the highest proportions of sequenced versus CoL-described species despite also being amongst the most species-rich clades: 0.063% sequenced species for Lepidoptera, 0.019% for Diptera, and 0.016% for Hymenoptera. An exception to this observation is Coleoptera -beetles, weevils, etc., which has the highest number of described species to date with currently available genome assembly resources for only 0.007% of these species.

These uneven distributions likely reflect historical biases in research interests for dipterans, which include the model species *Drosophila melanogaster* and disease-vectors like mosquitoes, for lepidopterans, which have been a model to study the genetic basis of complex traits and population genetics, and for hymenopterans, which include many well-studied social insects. While such biases may persist due to factors such as research priorities and ease of sampling, the balance should improve as the numbers and taxonomic spread of available arthropod genome assemblies continue to grow rapidly (Figure 1, Inset). Surveying taxonomic representation in this way highlights the increasingly rapid accumulation of new genome assemblies at the NCBI, providing researchers with a comprehensive overview of the species coverage of available genomics resources for their taxa of interest.

Assessing the surveyed species data allows for phylum-wide comparisons of the contiguity and completeness of genome assemblies available at the NCBI. Focusing on the 21 orders with at least five assemblies, order representation is notably unbalanced and assembly quality metrics summarised with N50 lengths and BUSCO completeness scores vary greatly among and within orders (Figure 2). Large differences between assembly and species counts are primarily driven in Lepidoptera by *Heliconius melpomene* (n=42), *Junonia neildi* (n=35), *Junonia evarete* (n=32), and six other *Junonia* and *Heliconius* species with more than 10 assemblies, and in Diptera mainly by *Drosophila melanogaster* (n=26), *Drosophila simulans* (n=12), and *Anopheles coluzzii* (n=10). The 307 species with more than one assembly comprise distinct assembly submissions and not updates that result in new versions of existing submissions (in this case only the latest version is surveyed). About half (142) of these species with multiple assemblies are represented by a chromosome-level assembly. Across all assemblies, those labelled as chromosome-level account for 12.3%, while a further 41.1% are labelled as scaffold-level assemblies, and the remaining 46.6% are contig-level (Figure S2).

**Figure 2.**
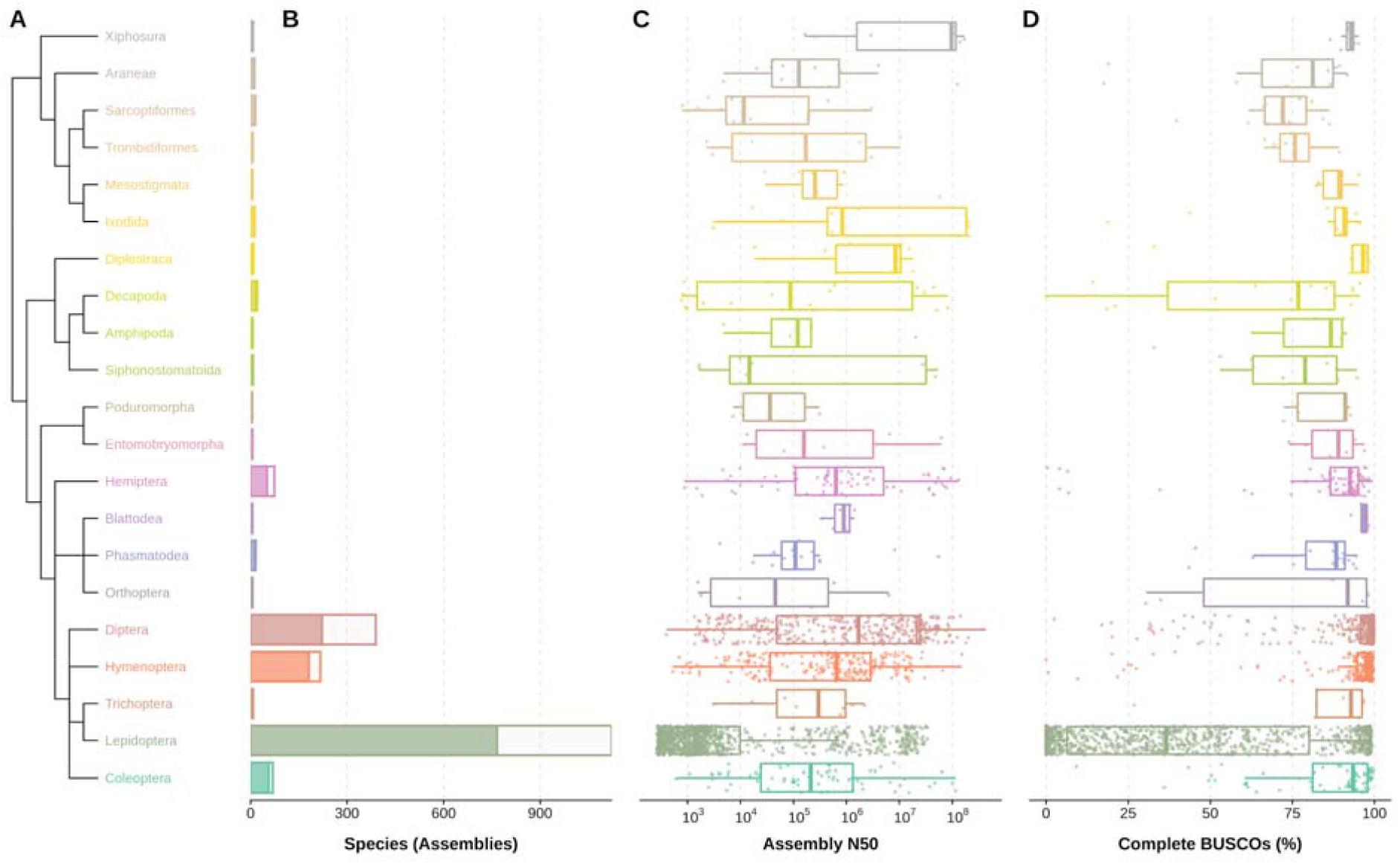
Order-level representation, contiguity, and completeness of 2024 available assemblies for 1326 arthropod species from the 21 orders with more than five assemblies. Data are presented only for orders with at least five assemblies available at the NCBI (2021-06-11). (A) Phylogenetic relationships of the 21 orders as resolved by the NCBI Taxonomy database. (B) Number of assemblies (entire bars) and unique species (dark fractions) retrieved from the NCBI Assembly database for each order. (C) Distribution of assembly NCBI scaffold N50 values (base pairs, log scale) for each order. (D) Distribution of Benchmarking Universal Single-Copy Orthologue (BUSCO) completeness (% of 1013 BUSCOs) for the arthropod lineage dataset (arthropoda_odb10) for each order. Boxplots show the median, first and third quartiles, and lower and upper extremes of the distribution (1.5 x IQR), and all values are overlaid as points to show the full distribution.

Excluding Lepidoptera that are skewed by a large number of poor-quality assemblies (Ellis et al. 2021), median N50 lengths per order represented by at least five assemblies (shown in Figure 2C) range from 11.6 Kbp for Sarcoptiformes (mites, 15 assemblies for 12 species) to 96.3 Mbp for Xiphosura (horseshoe crabs, 8 assemblies for 4 species). The horseshoe crabs have large genomes of 1.7-2.2 Gbp, for which concerted efforts have been successful in producing contiguous assemblies (Zhou et al. 2020; Shingate, Ravi, Prasad, Boon-Hui Tay, et al. 2020; Shingate, Ravi, Prasad, Boon□Hui Tay, et al. 2020; Nong et al. 2021). The mite genomes are all much smaller, with a median assembly span of just 88.5 Mbp, where the latest assembly for the parasitic mite, *Sarcoptes scabiei*, provides an example of how long-read technologies are helping to improve available genomic resources (Korhonen et al. 2020).

Median BUSCO completeness scores per order represented by at least five assemblies for the Arthropoda lineage dataset (Figure 2D) are less variable than the N50 lengths and, excluding Lepidoptera, range from 72.1% for Sarcoptiformes to above 97% for Diplostraca (clam shrimps and waterfleas, 9 assemblies for 7 species), Blattodea (cockroaches and termites, 6 assemblies for 5 species), Diptera, and Hymenoptera. Although within-order distributions can be highly variable, all but two of the 21 orders (Sarcoptiformes and Trombidiformes mites) are represented by at least one assembly with more than 90% complete BUSCOs. These contiguity and completeness distributions include all available assemblies, i.e. not filtered by level (contig, scaffold, chromosome) or type (haploid, principal or alternate pseudohaplotype, etc.). The completeness of contig-level assemblies is expectedly lower than that of scaffold-or chromosome-level (Figure S2B) assemblies, and although alternate pseudohaplotype assemblies can achieve high BUSCO completeness scores, they are generally lower than for principal pseudohaplotypes (Figure S2C). These phylum-wide comparisons of the qualities of available genome assemblies highlight the unbalanced order-level species representation as well as the variable levels of contiguity and completeness within and amongst arthropod orders.

### Arthropod assembly contiguity, size, and completeness

With 2083 assemblies exhibiting variable contiguities and sizes, the survey results provide the opportunity to examine expectations of how assembly contiguity and size relate to gene content completeness. Although long-read sequencing technologies are producing improved results (Hotaling et al. 2021), large genomes have often been challenging to assemble due to expanded proportions of repetitive sequences (Li et al. 2019). Even for smaller genomes, repeats can hinder scaffolding of contigs, reducing contiguity and possibly adding undetermined gap regions to the assembly. Less contiguous assemblies are thus expected to have more genes split across scaffolds, or partially or completely missing, resulting in lower completeness scores (Waterhouse et al. 2019).

The Earth BioGenome Project (Lewin et al. 2018) criteria for a reference quality assembly include obtaining a complete and single-copy BUSCO score above 90% and having the majority of sequences assigned to chromosomes. While 828 of the assessed arthropod assemblies achieve a complete and single-copy BUSCO score >90%, only 229 of these are also labelled as chromosome-level assemblies. Indeed, comparing assembly N50 values with their completeness scores shows that obtaining >90% complete BUSCOs can be achieved across a wide range of contiguities (Figure 3A). Recovery of >90% complete BUSCOs is observed for assemblies with N50s as low as 3.5 Kbp (*Tetragonula mellipes*, stingless bee, 92.1% Complete) and 3.9 Kbp (*Chrysomya rufifacies*, blow fly, 97.4% Complete). While some with N50s below 10 Kb are able to achieve >90% (n=25) or 80-90% (n=21) completeness, the vast majority of assemblies with such low contiguity levels achieve considerably lower BUSCO completeness scores than contiguous assemblies (i.e. N50 greater than 10 Kb). Among the latter, notable anomalies include 24 assemblies with N50s greater than 10 Kbp that nonetheless all have completeness scores of <50%. A third of these are labelled as alternate pseudohaplotypes, which offers an explanation for the low completeness levels as they likely represent collections of purged haplotigs. Others include improbably small assembly spans, e.g. *Sertania guttata* (butterfly, 30 Mbp span of 628 Mbp estimate) and *Dactylopius coccus* (scale insect, 18 Mbp span of 386 Mbp estimate), or high proportions of undetermined sequence, e.g. the brown recluse spider, *Loxosceles reclusa* (45% gaps). Biological complexity may also offer explanations, such as in the case of the Lord Howe Island stick insect, *Dryococelus australis* (N50=17.3 Kbp, 43.5% Complete), a potentially hexaploid genome with an estimated size of 4.2 Gbp that achieved an assembly span of 3.4 Gbp (Mikheyev et al. 2017).

**Figure 3.**
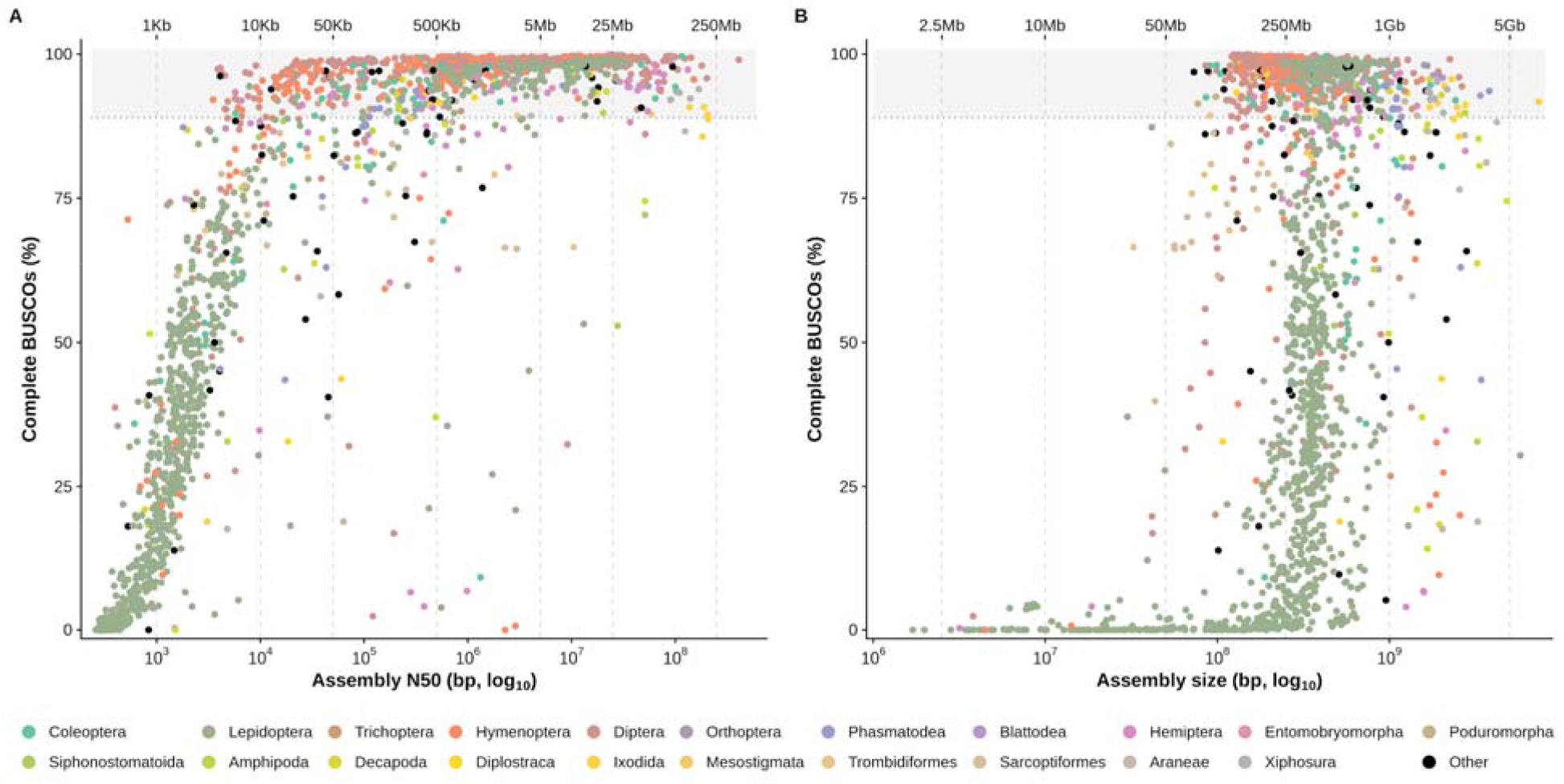
BUSCO completeness compared with assembly contiguity and size. Complete BUSCOs (in % of total BUSCOs for the arthropoda_odb10 dataset) are plotted against assembly N50 in bp (A) and assembly size in bp (B) for each assessed assembly. Both assembly N50 and assembly size are represented with a log scale. The color of a point indicates the order of the sequenced species. Dotted lines indicate N50 values of 1 Kbp, 10 Kbp, 50 Kbp, 500 Kbp, and 5 Mb in panel A and assembly size values of 50 Mb, 250 Mb, 1 Gb, and 5 Gb in panel B. BUSCO completeness scores above 90% are highlighted with a grey background.

The largest assemblies span more than 5 Gbp, with the maximum reported for the tick, *Haemaphysalis longicornis*, at 7.3 Gbp that shows 92% complete BUSCOs (Figure 3B). The estimated genome size for this tick however is only 3.4 Gbp, and a duplicated BUSCO score of 74.4% suggests that the applied assembly methods failed to collapse the alternative haplotypes. Indeed, an alternative assembly for this tick spans just 2.6 Gbp and scores 89.5% complete and 2.1% duplicated BUSCOs. A handful of other large assemblies with high duplicated scores are annotated as being non-collapsed, but others with many duplicated BUSCOs are also likely diploid or partially diploid (Figure S3). The smallest reported genome size for an arthropod to date is that of the tomato russet mite, *Aculops lycopersici* (Trombidiformes), exceptionally streamlined at only 32.5 Mbp (Greenhalgh et al. 2020). It achieves a Eukaryota completeness score of 83%, but only 67% Athropoda complete, which could reflect the evolutionary streamlining process but may also be related to challenges during gene prediction in such a gene-dense genome where genes have also experienced large-scale intron losses. The smallest assembly with a >80% Arthropoda completeness score is that of a grasshopper, *Xenocatantops brachycerus* (42 Mbp, 92% complete); however, inspecting the metadata reveals this to be a transcriptome rather than a genome assembly (Zhao et al. 2018). Amongst the smallest true genome assemblies achieving >80% completeness are other Trombidiformes as well as Sarcoptiformes, *e*.*g*. the house dust mite *Dermatophagoides farinae* (54 Mbp, 84% complete). Although there are fewer large assemblies spanning >1Gbp, across the full range of their sizes most achieve good completeness scores of >90%, indicating that sequencing technologies and assembly methods are able to overcome challenges often associated with large genomes.

Comparing assembly N50s and sizes with BUSCO duplicated scores (Figure S3) identifies several assemblies with high duplication levels. Some of these are labelled as ‘unresolved-diploid’ assemblies, which explains these high duplication levels, but this mechanism to inform users about the non-strictly-haploid status of certain assemblies is not widely nor consistently applied. Fragmented BUSCO scores (Figure S4) are expectedly higher for most of the less contiguous assemblies, highlighting those where many genes are likely split across two or more scaffolds. The survey results therefore provide the community with a comprehensive overview of genomic dataset qualities and of how contiguity and size relate to gene content completeness across currently available arthropod genome assemblies.

### BUSCO dataset lineage and version comparisons

The reference BUSCO lineage datasets are defined at different taxonomic levels that capture sets of near-universal single-copy orthologues from OrthoDB (Kriventseva et al. 2019) at ancient, intermediate, and younger nodes of the tree of life (Simão et al. 2015; Waterhouse et al. 2018). As duplication and loss events over evolutionary time erode the numbers of identifiable BUSCOs, datasets defined for more ancient lineages are smaller than for the younger ones, e.g. n=255 for Eukaryota and n=954 for Metazoa, versus n=3285 for Diptera and n=13780 for Primates (OrthoDB v10 datasets). An advantage of the smaller older lineage datasets is that compute runtimes are shorter because there are fewer individual genes to search for. The larger younger lineage datasets on the other hand offer greater resolution, meaning scores are impacted less by small differences in counts of complete, fragmented, or missing BUSCOs.

Our results provide the opportunity to compare the scores obtained using different lineage datasets for a large number of arthropod assemblies (Figure 4). Comparing percentages of complete BUSCOs identified with the Eukaryota (n=255) and the Arthropoda (n=1013) lineage datasets for a total of 1977 arthropod assemblies shows highly linearly correlated scores, especially for the highest-scoring assemblies (Figure 4A). For those scoring <80% there is a small but noticeable shift towards Arthropoda producing slightly higher scores than Eukaryota, indicating that proportionately more of the larger set of Arthropoda BUSCOs can be recovered from lower quality assemblies. Outlier points above the identity (y=x) axis suggest that the lower-resolution Eukaryota lineage dataset occasionally produces over-estimates of completeness, where proportionately more of the smaller set of ancient Eukaryota BUSCOs are recovered. Similar trends are observed when comparing the Arthropoda results to the higher resolution Insecta (n=1367) lineage dataset, with highly linearly correlated scores and occasional small over-estimates of completeness using the Arthropoda lineage dataset (Figure S5A).

**Figure 4.**
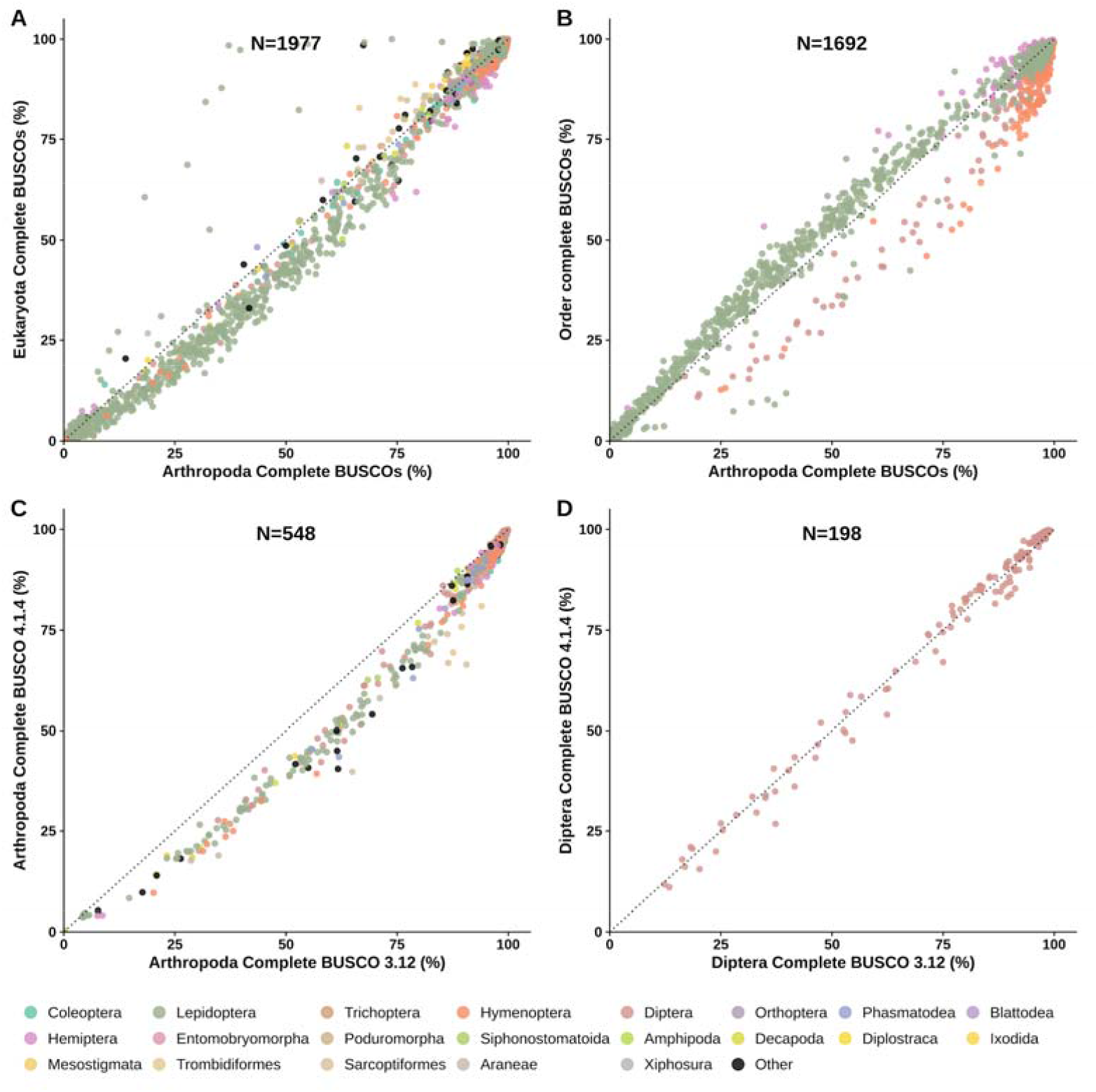
Comparisons of BUSCO lineage datasets and BUSCO versions. Congruence of BUSCO completeness scores are assessed by comparing results from (A) the Eukaryota (n=255) and the Arthropoda (n=1013) lineage datasets, (B) the Arthropoda and four insect order-level lineage datasets (Hemiptera n=2510; Hymenoptera n=5991; Lepidoptera n=5286; Diptera n=3285), and lineage datasets from BUSCO v4 (OrthoDBv10) and BUSCO v3 (OrthoDBv9) for (C) Arthropoda (odb9: n=1066) and (D) Diptera (odb9: n=2799). In each panel, the dotted lines show the identity (y=x).

Comparing Arthropoda results to those from four insect order-level lineage datasets shows high agreements for the highest-scoring assemblies (Figure 4B). For lower-scoring assemblies, results from applying the Lepidoptera and Hemiptera lineage datasets tend towards slightly higher scores than for Arthropoda. In contrast, using the Hymenoptera and Diptera lineage datasets generally produces lower completeness scores than for Arthropoda. These shifts could arise from the uneven representations of these orders in the 90-species Arthropoda lineage dataset which is dominated by 20 hymenopterans and 15 dipterans, with only 9 species each for Lepidoptera and Hemiptera. The same trends are observed when comparing results from the order-level lineage datasets to those from the Insecta dataset (Figure S5B).

In addition to updates to the codebase, BUSCO v4 was released with updated lineage datasets based on orthology data from OrthoDB v10 (Kriventseva et al. 2019), while BUSCO v3 used data from OrthoDB v9 (Zdobnov et al. 2017). Comparing completeness scores using the two Arthropoda datasets shows high levels of agreement for the highest-scoring assemblies with a consistent shift towards lower scores reported by BUSCO v4 for lower-quality assemblies (Figure 4C). A similar pattern is observed when comparing results from the two Insecta datasets (Figure S5C). The Diptera comparisons on the other hand reveal some score variations, which nevertheless agree well over the full range of assembly qualities (Figure 4D), similarly to results from the Hymenoptera datasets (Figure S5D). The different versions therefore produce generally consistent and comparable estimates of completeness, with a tendency for the OrthoDB-v10-based Arthropoda and Insecta datasets to report lower scores, especially for lower-quality assemblies. For objective quantitative comparisons it is thus necessary to assess assemblies using the same BUSCO versions, parameters, and lineage datasets, as presented here for the phylum-wide assessments of available arthropod genome assemblies.

### The Arthropoda assembly assessment catalogue: A^3^Cat

Running the workflow on the selected taxon of Arthropoda (NCBI:txid6656) produced the first version of the Arthropoda Assembly Assessment Catalogue (A^3^Cat v.2021-06-11), demonstrating how the workflow can be used to build a community resource. The A^3^Cat is provided as a searchable online table from https://rmwaterhouse.org/a3cat that allows browsing and downloading the collated metadata and BUSCO assessment results for arthropod assemblies available from the NCBI (n=2083 for A^3^Cat v.2021-06-11). Through simple text searches and/or applying query filters, users are able to quickly obtain downloadable overviews of the availability and quality of genome assembly resources for their arthropod taxa of interest. Without the computational burden of having to evaluate publicly available resources themselves, users can directly compare the assessments of their own assemblies with the precomputed results available from the A^3^Cat. In addition, for version and parameter controlled like-for-like comparisons, a user-workflow is provided to compute quality metrics on user-provided assemblies and compare them with A^3^Cat results for species from the same taxonomic clade (code and documentation are available from: https://gitlab.com/evogenlab/a3cat-user-workflow).

## Conclusions

Results from applying the assessment workflow to the phylum Arthropoda demonstrate the utility of building resources that provide a standardised overview of the current taxonomic coverage and quality of genome assembly resources available from the NCBI. The large-scale dataset also offers the opportunity to examine how widely used assembly metrics relate to BUSCO genespace completeness across a heterogeneous collection of genomes. Some anomalies point to errors or inconsistent use of metadata annotations where retractions or revisions would help to avoid misleading users about these resources. Furthermore, comparing results using different BUSCO datasets on large collections of assemblies reveals trends associated with employing ancient (lower resolution) or younger (higher resolution) lineages, and datasets built for BUSCO v3 or v4. While congruence is high especially for high-scoring assemblies, truly objective comparisons require reporting of the BUSCO versions, parameters, and lineage datasets used. Our data will enable future large-scale comparisons with results from the recently released BUSCO v5, which includes a new genome assessment strategy that improves efficiency and runtimes (Manni et al. 2021). The automated analysis workflow to build and maintain NCBI genome assembly assessment catalogues for selected taxa allows users to build updatable community resources, here exemplified with the A^3^Cat that facilitates surveying of species coverage and data quality for available arthropod assemblies and serves to guide ongoing and future genome generation initiatives.

## Materials and Methods

### Assembly selection and assessment workflow implementation

Accession numbers for all assemblies in the user-specified taxon are retrieved by querying the NCBI datasets application programming interface (API) (https://www.ncbi.nlm.nih.gov/datasets) with the ncbi-datasets-pylib library (version 12.3.0 in version 1.0 of a3cat-workflow) (Step 1 in Figure S1). For each assembly, the data package is downloaded to a temporary zip file using the *datasets* command-line utility (version 11.22.0 in version 1.0 of the a3cat-workflow). The nucleotide sequence and metadata are extracted from each data package with the ncbi-datasets-pylib library and stored as fasta and JSON files, respectively (Step 2 in Figure S1). For each assembly, complete taxonomic information is retrieved from the NCBI Taxonomy database (Schoch et al. 2020) using the ete3 python module (Huerta-Cepas et al. 2016), version 3.1.2 in version 1.0 of the a3cat-workflow) and stored in a JSON file (Step 3 in Figure S1). Taxonomic information is used to determine all BUSCO lineage datasets relevant for each assembly (Step 4 in Figure S1). During this step, assemblies are filtered by size, scaffold N50, and a manual filter list to discard assemblies which are too short and/or fragmented to contain any BUSCOs; this is necessary because BUSCO returns an error if no BUSCOs are found. The completeness of each assembly is assessed using BUSCO in genome mode and all other settings to default (Simão *et al*., 2015, version 4.1.4 in version 1.0 of the a3cat-workflow) for each applicable lineage dataset (Step 5 in Figure S1). The results folder generated by BUSCO is saved as a compressed archive with the exception of the BLAST database (blast_db) and BLAST input sequences (<run_name>/blast_output/sequences). The full results table, missing BUSCO list, and short summary are also retained in the final output for convenience. Metadata retrieved from NCBI and BUSCO scores for all assemblies are aggregated into a JSON file which summarises all the raw information retrieved and computed by the workflow (Step 6 in Figure S1). This JSON file is converted into a table with formatted headers stored in a tab-separated file where columns represent metadata and BUSCO scores and each line corresponds to an assembly (Step 7 in Figure S1). Finally, an interactive table is generated as an HTML page using the Data Tables JavaScript library (https://datatables.net, version 1.10.24 in version 1.0 of the a3cat-workflow) (Step 8 in Figure S1). The entire workflow is implemented using the Snakemake workflow management engine (Köster & Rahmann 2012; Mölder et al. 2021) and all software dependencies are managed by the Conda package manager; this implementation ensures that the workflow is portable and entirely reproducible. Parameters for each step of the workflow are specified in a YAML file and additional configuration files can be used to customize the table and HTML output. The code and documentation for the workflow are available from https://gitlab.com/evogenlab/a3cat-workflow.

### Assessment workflow deployment and data analyses

Results presented in this study were obtained by running version 1.0 of the a3cat-workflow on 2021-06-11. Species estimates were retrieved from the NCBI Taxonomy database using ete3 (version 3.1.2) on 2025-08-21 and from the Catalog of Life version 2021-06-10. Phylogenetic trees were automatically generated from NCBI taxonomy data with ete3. BUSCO scores for version 4.1.4 were obtained directly from the output of a3cat-workflow, while scores for version 3.12 were obtained with a development release version of the workflow available from https://gitlab.com/evogenlab/a3cat-workflow/-/releases/paper-busco-v3. Figures were generated with ggplot2 version 3.3.5 (Wickham 2016) and ggtree version 3.0.1 (Yu et al. 2017) in R version 4.1.0 (R Core Team 2021). All data-related figures, numbers, and supplementary material were generated with a Snakemake workflow (Mölder et al. 2021) available from https://gitlab.com/evogenlab/paper-a3cat using Snakemake version 6.3.0.

## Acknowledgments

This research was supported by Novartis Foundation for medical-biological research grant #18B116 and Swiss National Science Foundation grants PP00P3_170664 and PP00P3_202669 to RMW.

## Author Contributions

RMW conceived the study. RF developed the workflows and performed the analyses. RF and RMW wrote the manuscript, all authors read and approved the manuscript.

