## Supplementary Materials for "Assessing species coverage and assembly quality of rapidly accumulating sequenced genomes"

#### An automated workflow for assembly assessments

Reporting assembly contiguity and completeness is important because genome assemblies often form the basis of many downstream analyses. The reliability of any inferences or interpretations is influenced by these assessments, to greater or lesser extents depending on the context and research questions. Quality assessment summaries are therefore now a widespread requirement for publication of genomic resources, and databases hosting the data usually report contiguity statistics and sometimes also BUSCO summaries. However, provision of these metrics is sparse and dispersed, heterogeneous in format and execution (e.g. parameters, versions), not readily accessible at scale, and unable to keep pace with the rapidly accumulating data.

The automated workflow for assembly assessments allows users to systematically assess all assemblies available at the NCBI for a user-specified taxon of interest (Figure S1). The workflow is designed to perform on-demand updates to assess assemblies added to NCBI since the last run of the workflow, thereby providing up-to-date overviews of species coverage and data quality without having to re-assess previously examined datasets. The workflow steps are summarised in Figure S1 with the final output being a JSON file and a tabulated file containing all the information retrieved for each assembly, and an HTML / JavaScript table displaying the collated data. It is implemented using the Snakemake workflow management engine (Köster & Rahmann 2012; Mölder et al. 2021) with software dependencies managed by the Conda package manager. It is fully automated and can be configured using a yaml file to specify all options including the query for searching the NCBI Assembly database, BUSCO parameters, and the collated data to display in the output tables. The code and documentation are available from https://gitlab.com/evogenlab/a3cat-workflow.

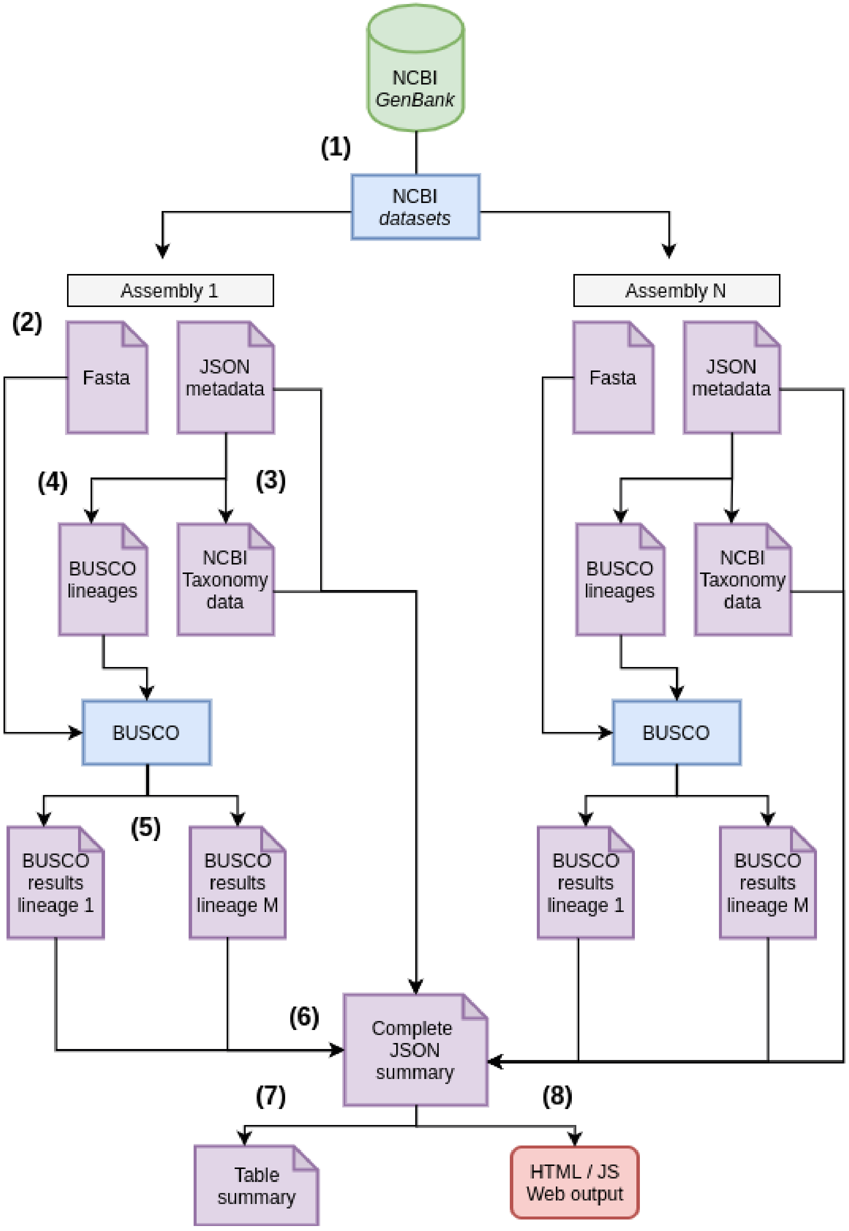


**Figure S1. Overview of the automated workflow for assembly assessments.**
The NCBI GenBank database is queried using the NCBI *datasets* python library **(1)** and assembly packages are downloaded with the *datasets* utility to obtain the genome sequence in a fasta file and metadata in a JSON file **(2)**. The complete taxonomy is retrieved from the NCBI taxonomy database for each assembly using ete3 **(3)** and used to determine relevant BUSCO lineage datasets **(4)**. BUSCO is then run with each lineage dataset on each assembly **(5)** and BUSCO results are aggregated with taxonomy information and metadata into a single complete JSON summary file **(6)**. Finally, the summary is converted to a tab-separated table **(7)** and an HTML / Javascript searchable table is generated **(8)**.

### Surveyed arthropod genome assembly resources

Assessing the surveyed species data allows for phylum-wide comparisons of the qualities of genome assemblies available at the NCBI. The NCBI Assembly database metadata labels assemblies as contig-level, scaffold-level, or chromosome-level, the majority of assemblies are labelled as scaffold-level (Figure S2A). While assemblies labelled as contig-level can achieve high BUSCO completeness scores, many achieve substantially lower completeness in contrast to the much higher completeness scores of most scaffold-level and chromosome-level assemblies (Figure S2B).

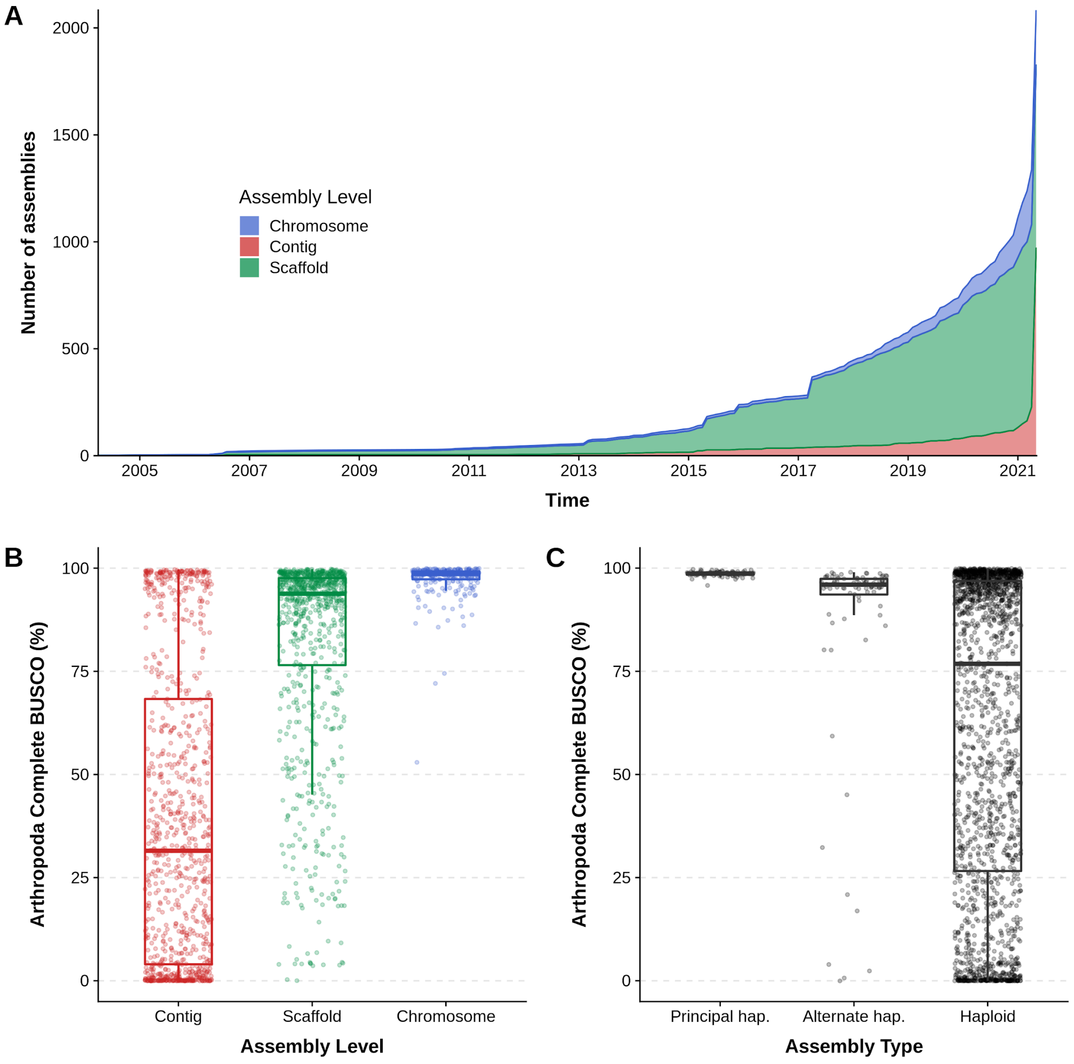

**Figure S2. Accumulation over time and BUSCO completeness of contig-level, scaffold-level, or chromosome-level assemblies.**
(**A**) The cumulative numbers of assemblies labelled as contig-level, scaffold-level, and chromosome-level according to their submission dates at the NCBI Assembly database. (**B**) Distributions of BUSCO completeness scores for assemblies labelled as contig-level, scaffold-level, and chromosome-level at the NCBI Assembly database, and (**C**) those labelled as simply haploid, or distinguishing between the principal and alternate haplotypes. Boxplots show the median, first and third quartiles, and lower and upper extremes of the distribution (1.5 x IQR), and all values are overlaid as points to show the full distribution.

### Contiguity, size, and BUSCO assessments of arthropod assemblies

Assemblies with the highest duplicated BUSCO scores (Figure S3) include those of *Drosophila elegans* (93.2%), *Aedes albopictus* (84.9%), *Haemaphysalis longicornis* (74.4%), *Anopheles maculatus* (54.8%), *Culicoides sonorensis* (50.1%), *Anopheles funestus* (50.1%), *Rhynchophorus ferrugineus* (40.4%), and *Anopheles coluzzii* (39.3%), none of which are annotated as being non-fully-collapsed assemblies. Others are labelled as 'unresolved-diploid' (another *Aedes albopictus* assembly, 64.4% duplicated) or 'haploid-with-alt-loci' (*Ixodes scapularis*, 39.5% duplicated). Thus while a mechanism exists to highlight to users the non-strictly-haploid status of assemblies it is not widely applied. The majority of assemblies with high levels of fragmented BUSCOs are themselves highly fragmented with low N50 values (Figure S4). Anomalies include the *Adela reaumurella* moth, with a scaffold N50 of 635 Kbp (874 Mbp assembly span composed of 610,001 scaffolds) and almost 40% fragmented BUSCOs, or the much more contiguous *Papilio bianor* butterfly (N50 13.1 Mbp) with nevertheless 26% fragmented BUSCOs. The large assembly of the *Locusta migratoria* locust (5.8 Gbp), with a completeness score of only 30%, shows a high fragmented BUSCO level of 30% suggesting that many genes are split across two or more scaffolds in this assembly.

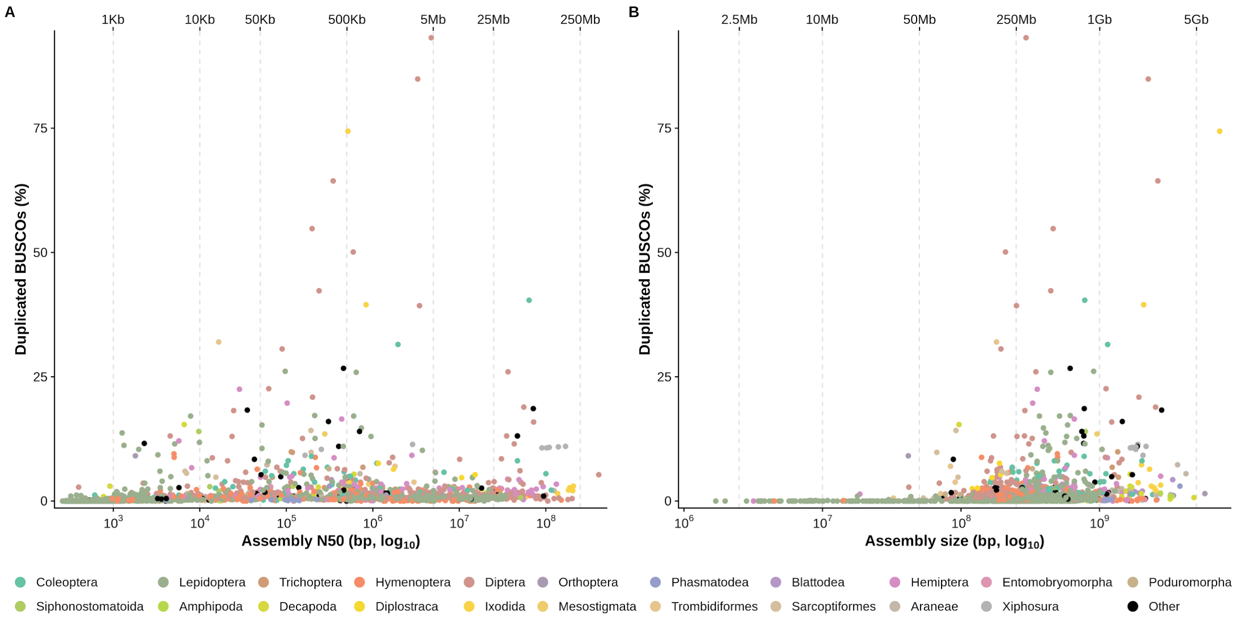


**Figure S3. Proportion of duplicated BUSCOs compared with assembly contiguity and size.** Duplicated BUSCOs (in % of total BUSCOs for the arthropoda_odb10 dataset) are plotted against assembly N50 in bp (**A**) and assembly size in bp (**B**) for each assessed assembly. Both assembly N50 and assembly size are represented with a log scale. The color of a point indicates the order of the sequenced species. Dotted lines indicate N50 values of 1 Kbp, 10 Kbp, 50 Kbp, 500 Kbp, and 5 Mb in panel **A** and assembly size values of 50 Mb, 250 Mb, 1 Gb, and 5 Gb in panel **B**.

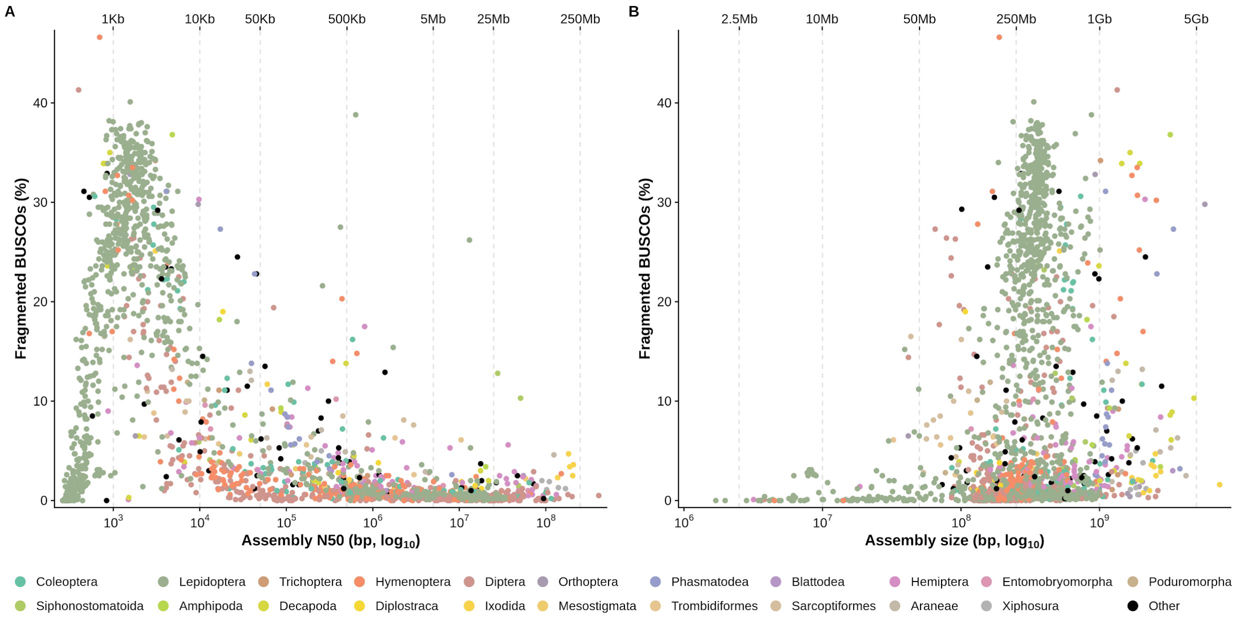


**Figure S4. Proportion of fragmented BUSCOs compared with assembly contiguity and size.** Fragmented BUSCOs (in % of total BUSCOs for the arthropoda_odb10 dataset) are plotted against assembly N50 in bp (**A**) and assembly size in bp (**B**) for each assessed assembly. Both assembly N50 and assembly size are represented with a log scale. The color of a point indicates the order of the sequenced species. Dotted lines indicate N50 values of 1 Kbp, 10 Kbp, 50 Kbp, 500 Kbp, and 5 Mb in panel **A** and assembly size values of 50 Mb, 250 Mb, 1 Gb, and 5 Gb in panel **B**.

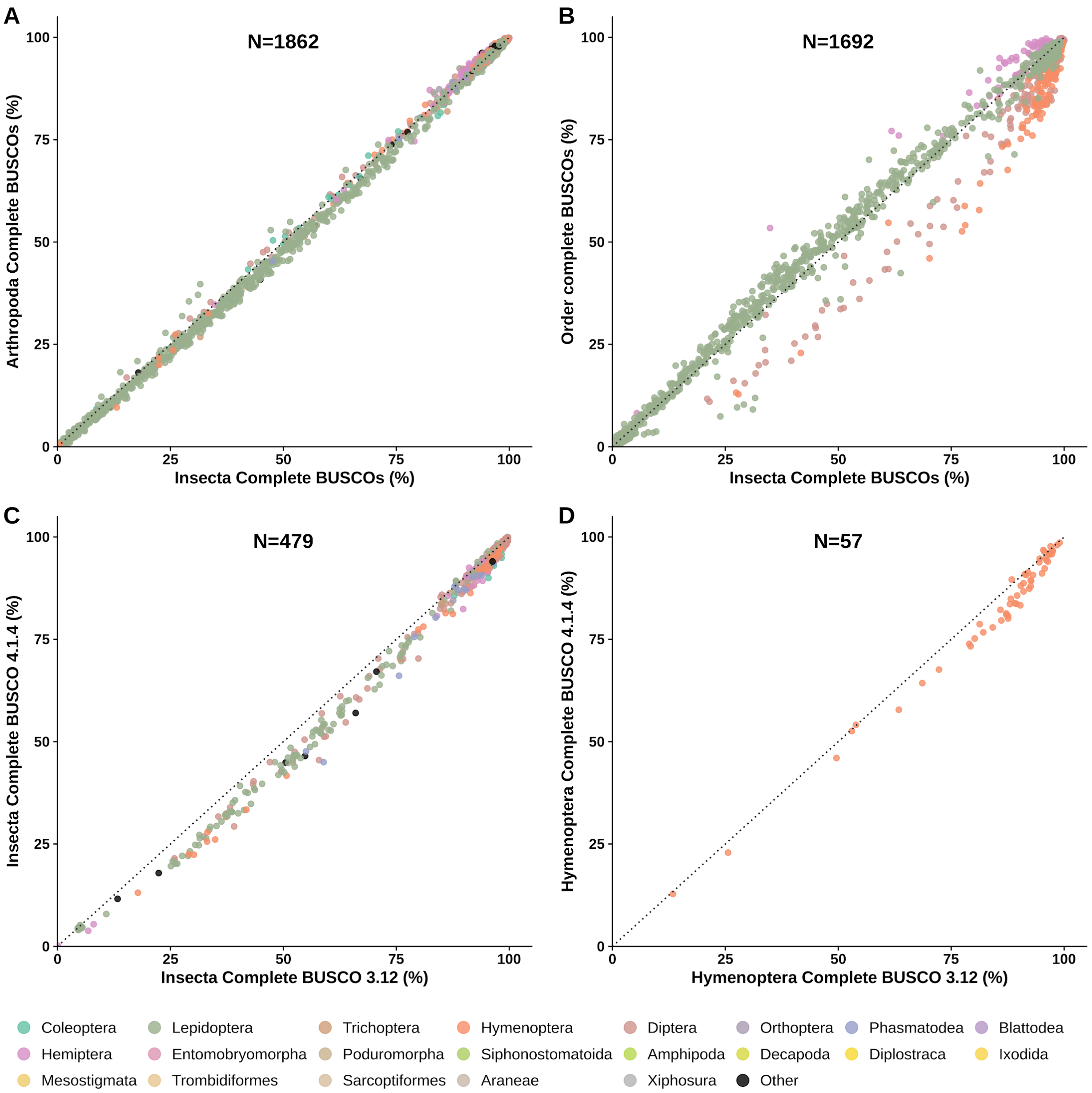


**Figure S5. BUSCO dataset comparisons for Insecta and Hymenoptera.**
Congruence of BUSCO completeness scores are assessed by comparing results from the Arthropoda (n=1013) and Insecta (n=1367) lineage datasets (**A**), the Insecta and four insect order-level lineage datasets (Hemiptera n=2510; Hymenoptera n=5991; Lepidoptera n=5286; Diptera n=3285) (**B**), and lineage datasets from BUSCO v4 (OrthoDBv10) and BUSCO v3 (OrthoDBv9) for Insecta (odb9: n=1658) (**C**) and Hymenoptera (odb9: n=4415) (**D**). Dotted lines represent the identity (y=x).
